# Sympathetic activation of sensory input and learning

**DOI:** 10.64898/2026.05.01.722216

**Authors:** Ellen Egeland Flø, Geir Magne Flø

## Abstract

A hallmark of learning is the need for sensory stimuli (Ginns, 2015; McGraw et al., 2009; Reinwein, 2012; Spence, 1950) so that learning is fundamentally based on sensory input signals affecting behaviour, physiology, and neurology. If behavioural measures of learning can be causally linked to physiological and neurological variables, a broader understanding of the mechanisms related to learning in schools, learning disabilities, and learning and health issues may emerge (McGraw et al., 2009). Despite decades of research on the physiological/neurological variable of sympathetic activation, learning, and achievement (Horvers et al., 2021), any causal relation remains unclear (Cowley et al., 2014; Mason et al., 2020; Pijeira-Díaz et al., 2016; Sung et al., 2023; Yu et al., 2024) and issues with instrument validation remain (Costantini et al., 2023; Hu et al., 2024; Milstein & Gordon, 2020; Van Der Mee et al., 2021). Here we investigate the effect of sensory input on sympathetic activation by using validated instruments for skin conductance measurement (Batista et al., 2019) and whether sympathetic activation is connected to learning in a cognitive laboratory context and an ecologically valid classroom context. In both contexts, we found a physiological variable which correlated with learning and that sensory input affected this variable while student movement did not. These sensory inputs varied depending on the different instructional activities the students participated in. Together, these findings bring us one step closer to a model linking sensory input to behavioural, physiological, and neurological variables.

---

Optimising the amount and quality of learning and teaching in schools remain a continuing concern for societies worldwide (Blikstad-Balas et al., 2022; Broughan et l., 2018; Harrison et al., 2022). This has led to a search for indicators of learning, in particular those that can be recorded in classroom settings and operate on timescales of minutes as opposed to whole lessons (Ahonen et al., 2018) to enable the assessment of different instructional activities regarding learning (Noroozi et al., 2020). Using physiological measurements as indicators of learning may provide insight into the biological processes that accompany it. Such an understanding would benefit learning technology development and learning intervention assessment through providing objective data not coloured by answering style, social desirability, or memory limitations (Cowley et al., 2014; Ravaja, 2004). Yet, no such mechanism has been proposed.

There has, however, been identified links between the physiological measure of skin conductance (SC) and learning (Cowley et al., 2014; Mason et al., 2020; Pijeira-Díaz et al., 2016; Sung et al., 2023; Yu et al., 2024), although the direction of the effect varies (Sung et al., 2023). Measurements of SC assess the sympathetic nervous system’s control of sweat gland secretion according to the tonic and phasic firing of sympathetically innervated eccrine nerves (Benedek & Kaernbach, 2010; Boucsein, 2012; Ellaway et al., 2010). Although skin conductance has been linked to learning, efforts to connect it to sympathetic activity in real-life settings are complicated by inconsistent signal-processing methods across studies (Horvers et al., 2021; Romine et al., 2022), for instance removing movement related variance in the SC signal. Such a practice may assume that cognitive activation causally affects sympathetic activation so that variance from other sources must be excluded from the SC signal when investigating cognitive activities such as learning. This assumption has still to be empirically validated.

An alternative mechanism behind this activation is that sensory input and movement affect sympathetic activation, which in turn affect learning through cognitive activation. This interpretation is supported by the improvement in working memory performance from listening to music, smelling perfume, and drinking coffee (Fekri Azgomi et al., 2023), and increased illuminance levels (Campbell et al., 2024). On a more fundamental level, basic sensory input from sight, hearing, touch, taste, and smell elicits SC responses (Bari et al., 2023; Gatti et al., 2018), indicating a link between sensory input and sympathetic activation, although SC responses were only compared across sensory input modalities and not to a baseline, limiting causal claims. Exercise, however, is known to activate the sympathetic nervous system (Christensen & Galbo, 1983; Lind, 1964; Martin et al., 1974) and increased physical activity improves learning (Petrigna et al., 2022; Sneck et al., 2019; Solberg et al., 2021).

Thus, to evaluate the hypothesis that sensory input and movement affect sympathetic activation, we report on two studies which investigated the effect of sensory input on SC in addition to the link between the intensity of sensory input, movement and sympathetic activation. To further strengthen/weaken the hypothesis that sympathetic activation is connected to learning, we assessed the relationship between SC measures and learning, and whether instructional activities affected sensory input, movement, and SC.

## Sympathetic activation and learning

Here we report one classroom study (study one) and one cognitive laboratory study (study two) to satisfy both relevance for real-life settings and causality claims. In the classroom study 12 male students (17– 18 years) participated in a physics education lesson whilst SC and video data was collected. Later, the students took a post-test consisting of 40 multiple choice questions (see ‘Methods – Participants and context – Study one’). In the lab study, 100 participants (59 female, 39 male, two preferred not to disclose) aged 12-21 years, without any self-reported learning disabilities or psychiatric diagnoses, were included. They answered the Spatial Reasoning Instrument (Ramful et al., 2017) to assess spatial ability which is closely related to learning in STEM (science, technology, engineering, and mathematics) (Wai et al., 2009) and a pre-/post-test with 10 multiple choice questions in biology. SC data was collected as the participants watched a six-minute instructional video in biology shown randomly in two versions with different intensity levels of sensory stimuli (see ‘Methods – Participants and context – Study two’).

To investigate the role of sensory input on sympathetic activation and learning, we deployed custom software to calculate the three sensory input variables, brightness, normalised brightness change, and loudness as defined in Figure 1. To include any role of student movement on sympathetic activation and learning, we used the multi-person human pose detection library OpenPose (Cao et al., 2021) to detect student poses with a combination of available software (Hur & Bosch, 2022) to track students across video frames and custom software to calculate movement from the coordinates as defined in Figure 1. This novel approach allowed simultaneous recording of sensory input and movement calculated from video data to be synchronised with SC measurements in both lab and freely moving classroom settings. The measurement and analysis of SC is shown in Figure 1 and ‘Methods – SC measurement and analysis’.

**Fig. 1.**
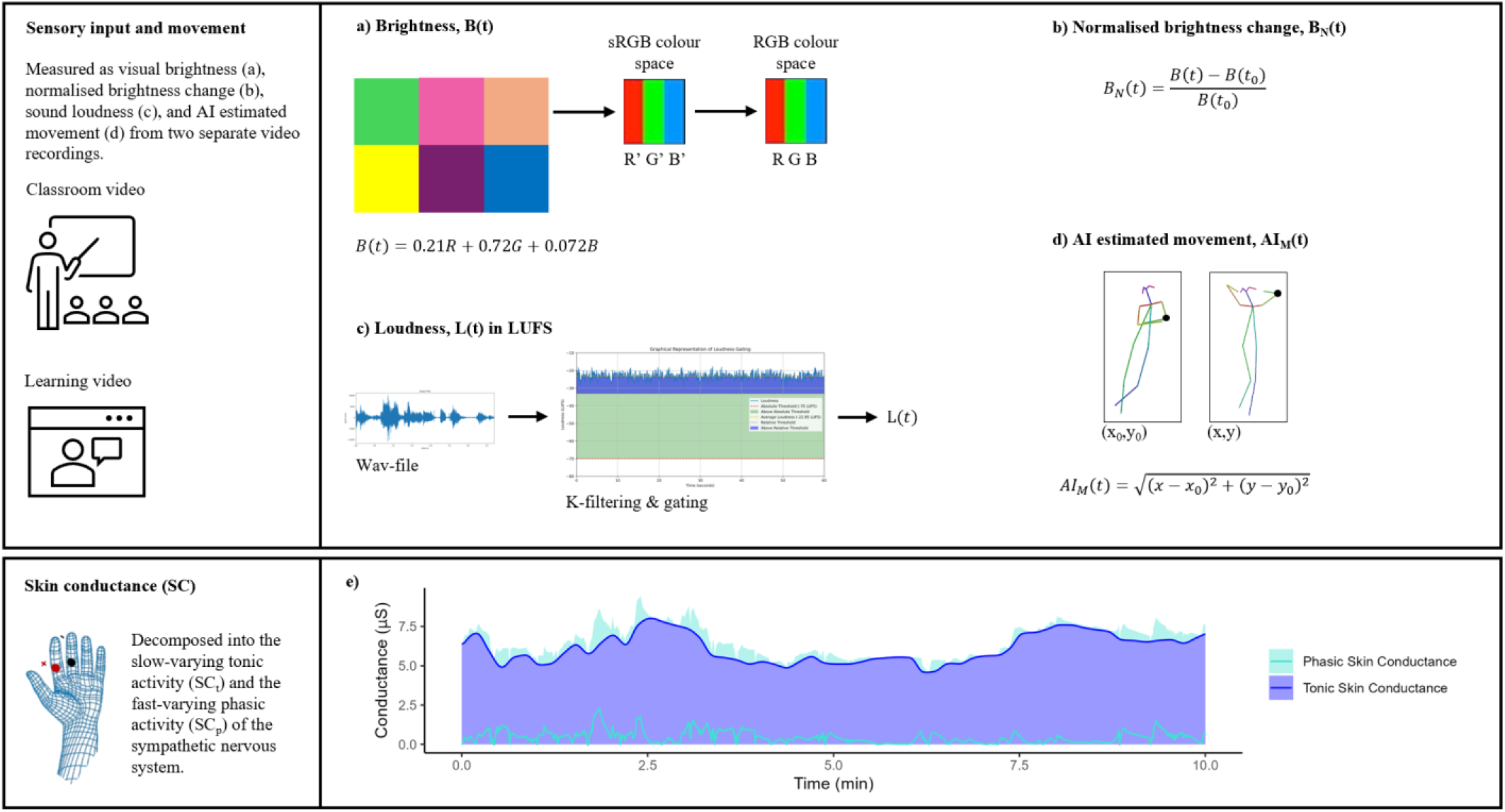
Analysing sensory input, movement, and skin conductance. **a**, Video recordings were used to assess brightness through transforming the sRGB colour space values of the video to the RGB colour space before linear addition according to human sensitivity to different light wavelengths was performed (see Methods - Video analysis - Brightness). **b**, We define normalised brightness change to be the difference in brightness between two consecutive frames of video relative to the brightness of the first frame. **c**, Using our software, we assessed loudness by analysing the wav-file from the video into LUFS values (LUFS, i.e., Loudness Units relative to Full Scale is a common unit for measuring sound intensity as experienced by human perception (International Electrotechnical Commission, 2003)) (see Methods - Video analysis - Loudness). **d**, By applying the multi-person human pose detection library (https://github.com/CMU-Perceptual-Computing-Lab/openpose) from OpenPose, we identified the student poses in study 1 and used a combination of available software (Hur & Bosch, 2022) for tracking students across video frames with our own software operationalising our definition of the AI estimated movement, namely, the cartesian distance between model identified body parts in consecutive video frames. **e**, The skin conductance was decomposed into the fast-varying phasic component and the more slowly varying tonic component by LedaLab software (version 3.4.9; http://www.ledalab.de/) for MATLAB using continuous decomposition analysis (CDA) (Benedek & Kaernbach, 2010) and the phasic component was used in further analysis as it has been connected to cognitive activation (Pijeira-Díaz et al., 2018) (see Methods - SC measurement and analysis).

To assess whether sympathetic activation was connected to learning, we measured SC while students were learning physics in the classroom study and checked for a correlation with the post-test. Here, neither tonic nor phasic mean (M) or standard deviation (SD) SC correlated significantly with learning in class. However, as both phasic SC measures approached significance whilst being highly correlated with a risk of multicollinearity issues (Lavery et al., 2019), we chose to combine them into a composite variable, SC_Phasic_M_/SC_Phasic_SD_, to minimise risk of type 2 error (see ‘Methods - Statistical analyses - Correlation and multiple regressions’ for more details). Thus, the composite phasic sympathetic activation, SC_Phasic_M_/SC_Phasic_SD_, correlated with test scores in the classroom study (r(10) = -0.71, P =.01, 95% CI = [-0.91, -0.22]). To further validate this finding in another context, namely the lab study, we measured SC while students were watching a biology learning video and checked for a correlation with the difference in pretest and post-test scores. Similarly, we found that SC_Phasic_M_/SC_Phasic_SD_ correlated with the difference in post- and pre-test scores in the laboratory study (β = -0.19, P = 0.04, 95% CI = [-0.38, -0.006]) when corrected for gender (male = -1, female = 1) (β = -0.16, P = 0.10, 95% CI = [-0.35, 0.03]) and spatial ability (β = 0.40, P = 3.1×10^-5^, 95% CI = [0.22, 0.59]) in the model (Pratt adjusted R^2^ = 0.18, F(3,94) = 8.45, P = 5.0×10^-5^) as shown in Figure 2.

**Fig. 2.**
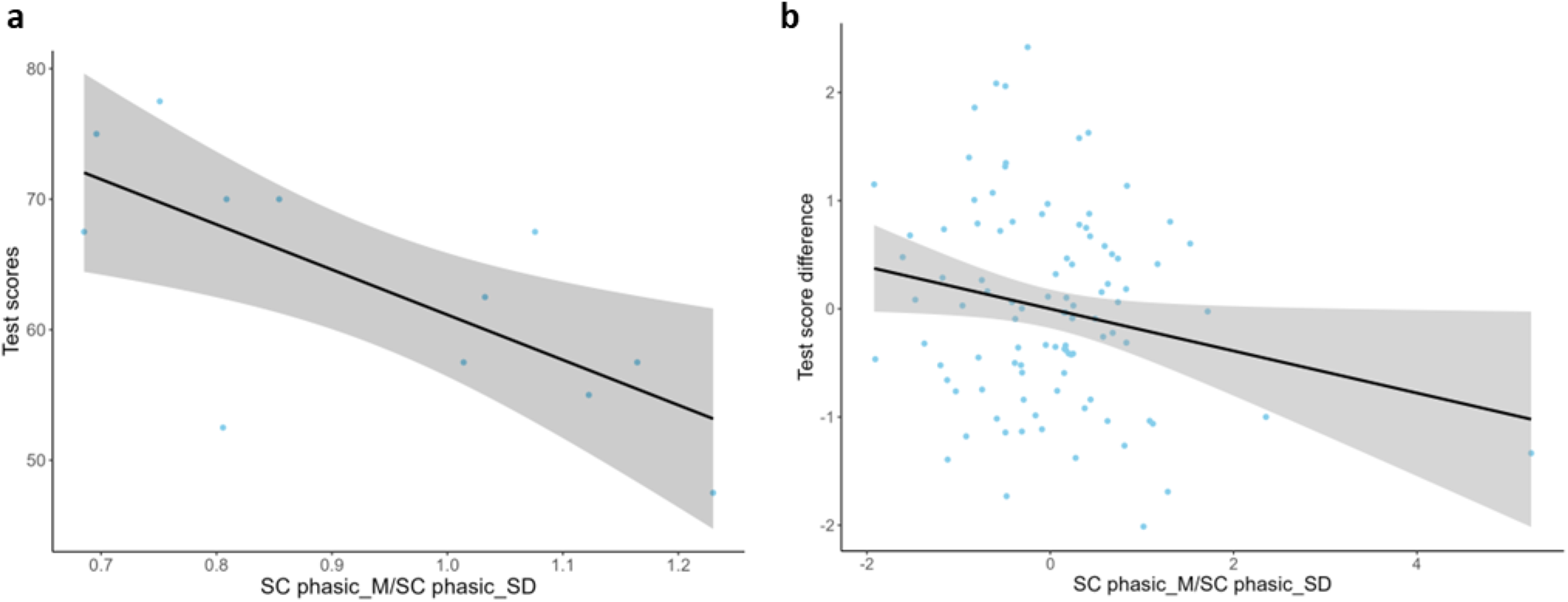
The connection between composite phasic sympathetic activation and learning. **a**, The Pearson correlation of the SC_Phasic_M_/SC_Phasic_SD_ and the test scores for n = 12 in study one. **b**, The Pearson correlation of the SC_Phasic_M_/SC_Phasic_SD_ and for learning as measured by the difference in post-test scores and pretest scores for n = 98 in study two in a linear multiple regression corrected for gender and spatial ability with normalised scores. The modelling assumptions held for the SCPhasic_M/SCPhasic_SD, test score variables, gender, and spatial ability (see ‘Methods - Statistical analyses - Correlation and multiple regressions’).

That the correlation is negative does not mean that activation is negatively connected to learning, as SC_Phasic_M_/SC_Phasic_SD_ is a composite variable with non-linear response to increased phasic activation. Because the composite phasic SC variable, SC_Phasic_M_/SC_Phasic_SD_, have high values for both very low (i.e., very low mean and very low SD) and very high (i.e., very high mean and medium SD) phasic sympathetic activation and low values for medium phasic sympathetic activation (i.e., medium mean and high SD), this finding aligns with findings of an optimal level of activation for performance in the dancing mouse (Yerkes & Dodson, 1908).

## Sensory input, movement, and sympathetic activation

We further wished to test whether sensory input affected sympathetic activation by showing one of two versions of a six-minute learning video to 98 students in the lab in study two. Video one had elevated saturation and slightly elevated volume compared to video two. To assess whether the videos demonstrated significant differences in sensory input, the three sensory input variables were compared for each video with the paired Wilcoxon signed-rank test and the corresponding effect size, rank biserial correlation, r. Brightness was lower for video one than for video two (W = 428, P = 0.03, r = 0.57, 95% CI = [0.30, 0.79]), whilst normalised brightness change was higher for video one than for video two (W= 981, P = 6.9×10^-6^, r = 0.87, 95% CI = [0.87, 0.87]), as was loudness (W = 1225, P = 2.2×10^-16^, r =0.87, 95% CI = [0.87, 0.87]). Did these sensory input differences affect sympathetic activation? Yes. We compared the composite phasic sympathetic activation for the students who viewed video one and video two and the SC_Phasic_M_/SC_Phasic_SD_ was higher for the video one group with the Wilcoxon rank sum test (W = 864, P = 0.03, r = 0.22, 95% CI = [0.04, 0.41]). Similar results were found for tonic sympathetic activation (W = 83, P = 3.1×10^-12^, r = 0.74, 95% CI = [0.63, 0.83]), whereas phasic activation demonstrated a significant difference in the opposite direction (W = 1171, P = 4.5×10^-14^, r = 0.78, 95% CI = [0.69, 0.84]) (See Figure 3, panels a-d). Thus, basic sensory input affect tonic, phasic, and composite sympathetic activation when the content of the sensory input is controlled for.

**Fig. 3.**
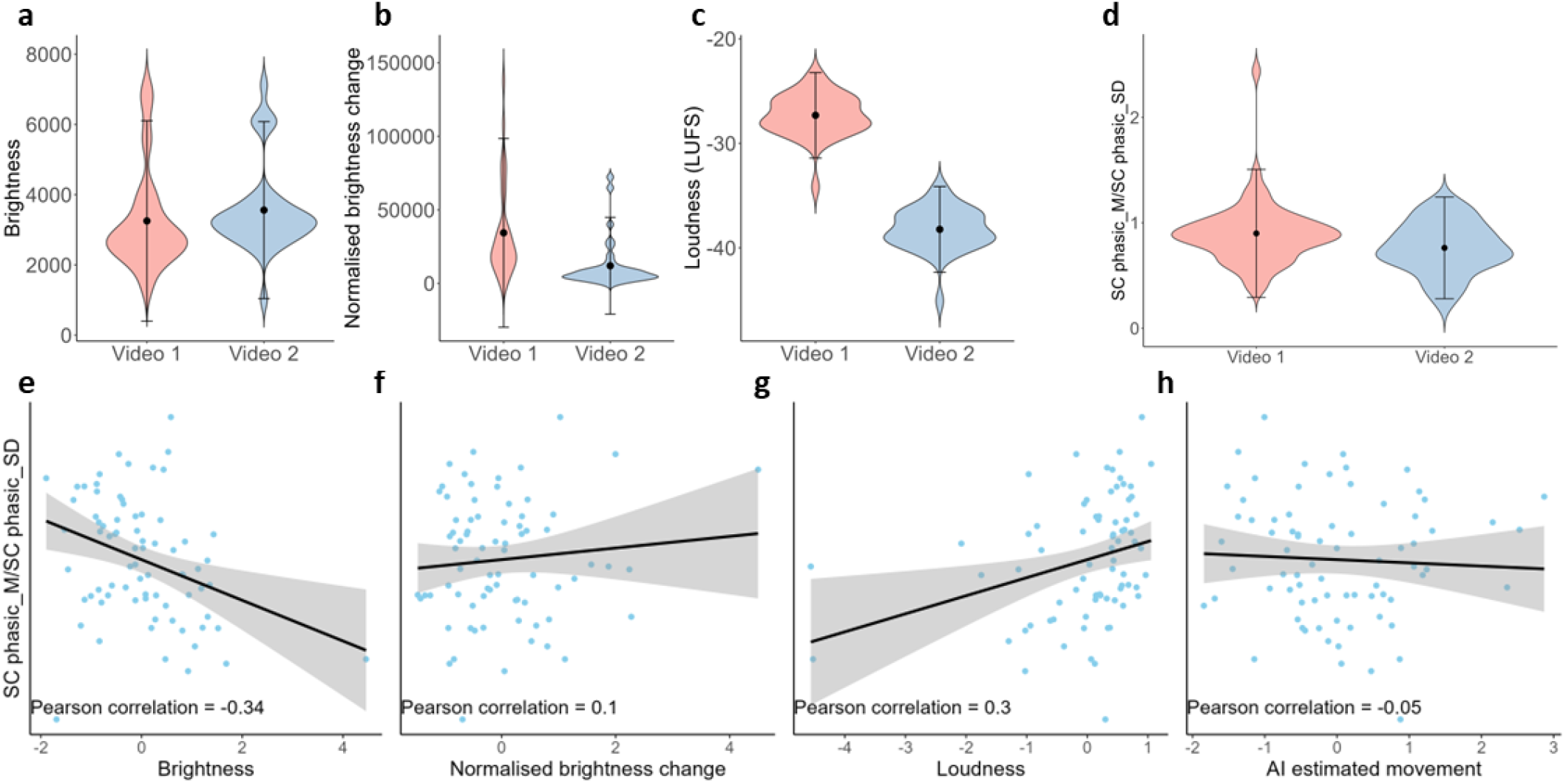
The effect of sensory input and movement on composite phasic sympathetic activation. **a**, The brightness levels of video 1 and video 2. **b**, The normalised brightness change levels of video 1 and video 2. **c**, The loudness levels in LUFS of video 1 and video 2. **d**, The SC_Phasic_M_/SC_Phasic_SD_ variable for the two sensory input conditions. The difference in the SC variable was still significant after removing the outlier in Video 1. **e**, The Pearson correlation of the SC_Phasic_M_/SC_Phasic_SD_ and brightness in study one in a linear multiple regression corrected for the other sensory input and movement variables, with normalised scores. **f**, The Pearson correlation of the SC_Phasic_M_/SC_Phasic_SD_ and normalised brightness change in study one in a linear multiple regression corrected for the other sensory input and movement variables, with normalised scores. **g**, The Pearson correlation of the SC_Phasic_M_/SC_Phasic_SD_ and loudness in study one in a linear multiple regression corrected for the other sensory input and movement variables, with normalised scores. **h**, The Pearson correlation of the SC_Phasic_M_/SC_Phasic_SD_ and AI estimated movement in study one in a linear multiple regression corrected for the sensory input variables, with normalised scores. The modelling assumptions held for the SC_Phasic_M_/SC_Phasic_SD_, test score variables, gender, and spatial variability (see ‘Methods - Statistical analyses - Correlation and multiple regressions’).

To further investigate the relationship between sensory input, movement, and composite phasic sympathetic activation in a more naturalistic setting, we estimated a linear multiple regression model with the sensory input variables as well as the AI estimated movement for the 75-minute lesson for the 12 students in the classroom in study one. Brightness predicted composite phasic sympathetic activation (β = -0.36, P = 0.001, 95% CI = [-0.57, -0.15]), normalised brightness change predicted sympathetic activation (β = 0.24, P = 0.04, 95% CI = [0.01, 0.46]), loudness predicted sympathetic activation (β = 0.35, P = 0.002, 95% CI = [0.13, 0.57]), whereas AI estimated movement did not predict sympathetic activation (β = -0.14, P = 0.22, 95% CI = [-0.37, 0.08]) in the model (Wherry adjusted R^2^ = 0.20, F(4,70) = 5.96, P = 0.0003) as shown in Figure 3, panels e-h. Tonic SC was predicted by normalised brightness change and loudness, whereas phasic SC was predicted by brightness, normalised brightness change, and loudness (See Extended Data Tables 2 and 3). This means that sensory input, but not movement, can predict sympathetic activation without controlling for the sympathetic activation caused by the students’ production of talk. That movement did not affect the sympathetic activation differs from previous research on the effect of exercise on sympathetic activation (Christensen & Galbo, 1983; Lind, 1964; Martin et al., 1974).

## Instructional activities

In study one, the 12 students participated in a physics lesson where the instructional activities included five types of activities. The 75 one-minute segments of video were assigned the dominant category and the composite phasic sympathetic activation, SC_Phasic_M_/SC_Phasic_SD_ was calculated for each activity type (see the ‘Video analysis – Instructional methods’ section of the Methods for more details). Namely, the teacher demonstrating content on the blackboard without student discussion (M = 19.32, SD = 39.75), the teacher demonstrating content on the blackboard with student questions and discussion (M = 3.14, SD = 8.05), teacher-led discussion without using the blackboard (M = 7.81, SD = 17.96), students working on physics tasks without teacher help (M = 13.98, SD = 28.61), and students working on physics tasks with teacher help (M =9.43, SD = 19.07). We further investigated the effect of type of instructional activity on composite phasic sympathetic activation for all pairwise combinations of instructional activities with the non-parametric Mann-Whitney U test for group comparison as our data was not paired with the corresponding effect size, the rank biserial correlation, r.

The blackboard with questions activity demonstrated lower SC_Phasic_M_/SC_Phasic_SD_ values than teacher-led discussion (V = 447931, P = 2.2×10^-16^, r = 0.01, 95% CI = [0.002, 0,13]), physics tasks with help (V = 605550, P = 2.2×10^-16^, r = 0.07, 95% CI = [0.006, 0.16]), physics tasks without help (V = 349866, P = 2.2×10^-16^, r = 0.16, 95% CI = [0.06, 0.27]), and blackboard without questions (V = 296835, P = 2.2×10^-16^, r = 0.16, 95% CI = [0.04, 0.28]). Similarly, teacher-led discussion demonstrated lower SC_Phasic_M_/SC_Phasic_SD_ values than physics tasks with help (V = 203841, P = 2.2×10^-16^, r = 0.08, 95% CI = [0.006, 0.20]), physics tasks without help (V = 70125, P = 2.2×10^-16^, r = 0.18, 95% CI = [0.04, 0.31]), and blackboard without questions (V = 47586, P = 2.2×10^-16^, r = 0.22, 95% CI = [0.07, 0.36]). Physics task with help also demonstrated lower SC_Phasic_M_/SC_Phasic_SD_ values than physics tasks without help (V = 139656, P = 2.2×10^-16^, r = 0.11, 95% CI = [0.007, 0.22]), and blackboard without questions (V = 106953, P = 2.2×10^-16^, r = 0.13, 95% CI = [0.02, 0.03]). Lastly, physics tasks without help demonstrated lower SC_Phasic_M_/SC_Phasic_SD_ values than blackboard without questions (V = 19701, P = 2.2×10^-16^, r = 0.08, 95% CI = [0.005, 0.27]), as shown in Figure 4. Because instructional activities were varied across the lesson, the composite phasic sympathetic activation differences indicate an effect of the specific classroom activities on students’ composite phasic sympathetic activation.

**Fig. 4.**
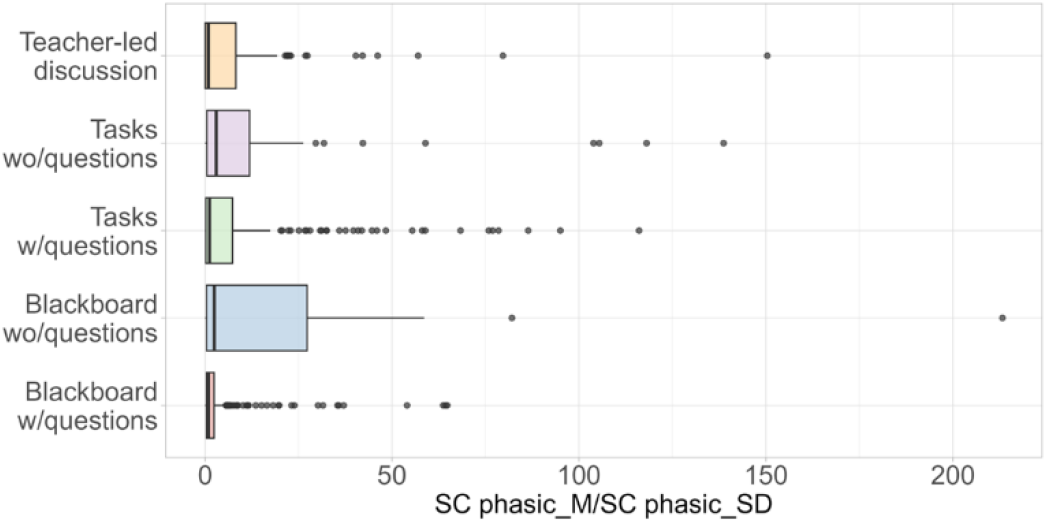
The effect of instructional activities on composite phasic sympathetic activation. The distribution of SC_Phasic_M_/SC_Phasic_SD_ measurements for the five different instructional activities.

Continuing to investigate instructional activities, we assessed their effect on sensory input and movement, to check whether the effect of instructional activities on composite phasic sympathetic activation may be mediated by their effect on sensory input. We compared brightness levels, normalised brightness change levels, loudness, and AI estimated movement per second for all pairwise combinations of instructional activities with the non-parametric Mann-Whitney U test for group comparison as our data was not paired with the corresponding effect size, the rank biserial correlation, r.

Brightness was significantly different for seven out of ten pairwise combinations starting with tasks without help being the lowest (M = 639.03, SD = 23.97), and in increasing order; teacher-led discussion (M = 640.63, SD = 17.43), blackboard without questions (M = 642.88, SD = 37.53), blackboard with questions (M = 644.57, SD = 25.87), and tasks with help (M = 644.76, SD = 16.36).

The significant effect sizes ranged from r = 0.05 to 0.21 (significant p-values = [0.01, 9.4×10^-7^]). See Extended Data Tables 4 and 5 for more details. For normalised brightness change, mainly a measure of change in visual sensory input but including some movement variance, the increasing order of the instructional activities were as follows, blackboard without questions (M = 25.16, SD = 12.38), blackboard with questions (M = 28.07, SD = 14.41), teacher-led discussion (M = 30.09, SD = 10.76), tasks with help (M = 43.94, SD = 16.50), and tasks without help (M = 44.67, SD = 16.13), with all but one group comparison being significant. The significant effect sizes ranged from r = 0.07 to 0.54 (significant p-values = [0.0003, 2.2×10^-16^]). See Extended Data Tables 6 and 7 for more details.

Loudness, mainly a measure of auditory sensory input but including some auditory output variance, similarly demonstrated significant differences for eight out of ten comparisons with effect sizes ranging from r = 0.09 to 0.49 (significant p-values = [1.2×10^-5^, 2.2×10^-16^]). The increasing order of the activities were blackboard without questions (M = 307.13, SD = 262.88), blackboard with questions (M = 509.45, SD = 1200.94), teacher-led discussion (M = 542.61, SD = 623.32), tasks without help (M = 653.10, SD = 569.69), and tasks with help (M = 820.45, SD = 800.59). See Extended Data Tables 8 and 9 for more details.

Lastly, the AI estimated movement was significantly different for seven out of ten pairwise combinations where the significant effect sizes ranged from r = 0.06 to 0.18 (significant p-values = [0.03, 2.2×10^-16^]). The increasing order of the activities were blackboard without questions (M = 394.37, SD = 287.61), teacher-led discussion (M = 560.99, SD = 717.94), blackboard with questions (M = 571.71, SD = 743.83), tasks with help (M = 647.44, SD = 747.25), and tasks without help (M = 656.89, SD = 863.87), following a similar pattern to normalised brightness change. See Extended Data Tables 10 and 11 for more details. These results are shown in Figure 5, panels a-d, and indicate that instructional activities have an effect on sensory input and movement (brightness, normalised brightness change, loudness, and AI estimated movement). Thus, sympathetic activation during instructional activities seems to occur alongside changes in sensory input and movement, suggesting these factors are closely bound together.

**Fig. 5.**
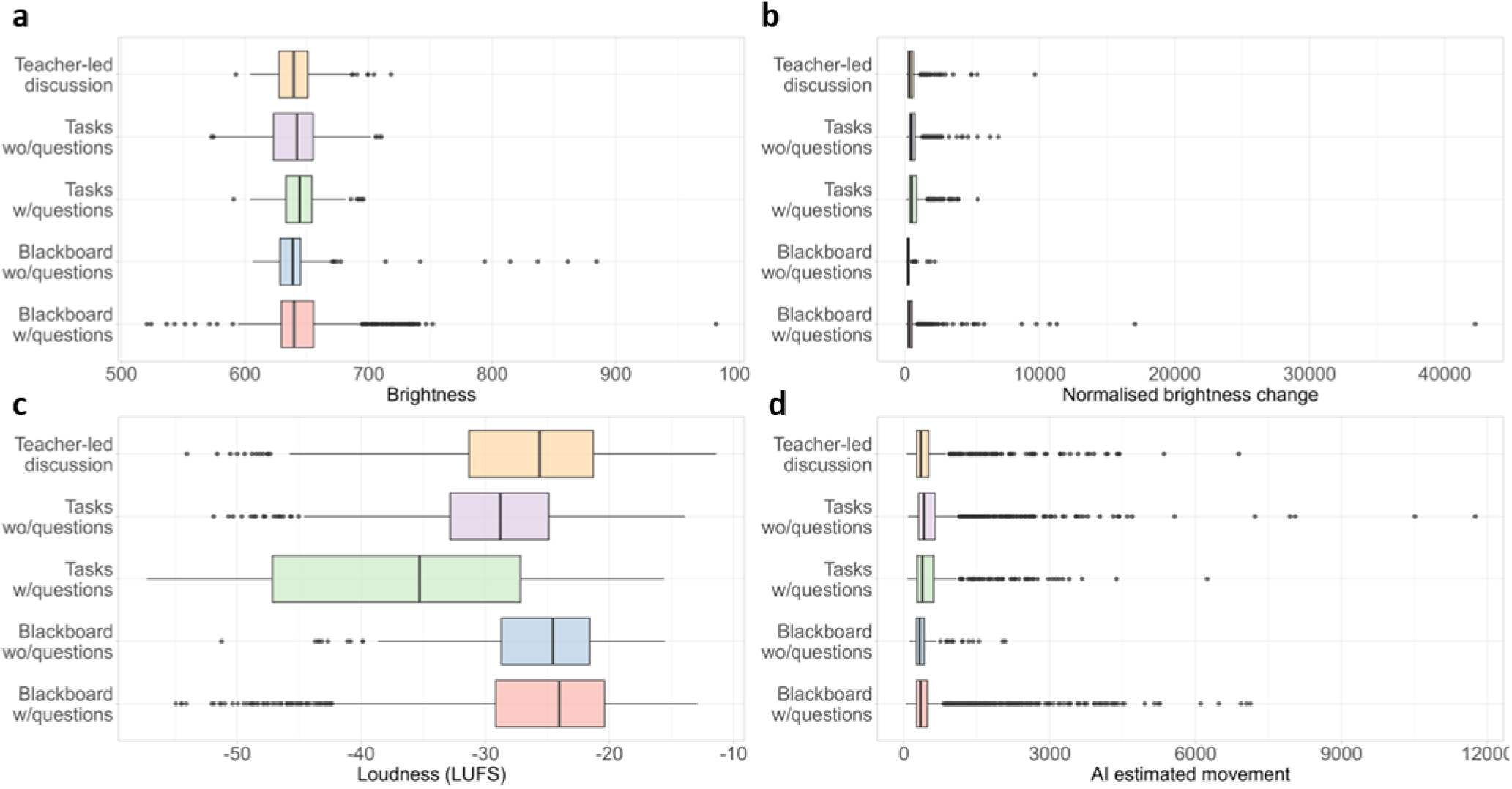
The effect of instructional activities on sensory input and movement. **a**, The distribution of brightness measurements for the five different instructional activities. **b**, The distribution of normalised brightness change measurements for the five different instructional activities. **c**, The distribution of loudness measurements for the five different instructional activities. **d**, The distribution of AI estimated movement measurements for the five different instructional activities.

## Discussion

Both included studies replicate the finding that a composite phasic SC variable negatively correlates with learning and that this variable is affected by basic sensory input in addition to different instructional activities. This pattern is consistent across two different contexts, namely, a cognitive laboratory and a naturalistic classroom setting. These findings support the suggested mechanism that sensory input affects sympathetic activation (as measured by SC) and that this sympathetic activation affects learning. These findings add to the limited research on sympathetic activation and learning (Cowley et al., 2014; Mason et al., 2020; Pijeira-Díaz et al., 2016; Sung et al., 2023; Yu et al., 2024) and align with previous studies on sensory input and sympathetic activation which found improved working memory performance and elicitations of SC resulting from sensory input across modalities (Campbell et al., 2024; Bari et al., 2023; Fekri Azgomi et al., 2023; Gatti et al., 2018).

The finding that sensory input affects sympathetic activation carries implications for the causal interpretation of this response, which differs from that assumed in previous research (Ikehara & Crosby, 2005; Nourbakhs et al, 2012; Setz et al., 2010; Shi et al., 2007). In cognitive load research, sympathetic activation is typically treated as an effect of cognitive load (Larmuseau et al., 2019). Consequently, parts of the SC signal are removed (e.g., that which corresponds to faster breathing, emotions, mental fatigue, and stress (Boffet, et al., 2025; Buchwald et al., 2019; Setz, et al., 2010)), before estimating cognitive load. Yet if these same factors trigger sympathetic activation which subsequently affects cognitive load and learning, such editing is unnecessary. It would imply that variations in the SC signal reflect the same underlying physiology regardless of whether their origin is cognitive or sensory.

However, although cognitive activity has been found to affect sympathetic activation (Ikehara & Crosby, 2005; Nourbakhs et al, 2012; Shi et al., 2007), such effects may reflect only the content of sensory input when basic sensory input is uncontrolled. This highlights the need for future research which accounts for basic sensory input to clarify whether the underlying causal pathways are bidirectional.

In naturalistic settings, controlling variables is challenging (Bronfenbrenner, 1977) and our instructional activities in the classroom study were naturally occurring across the lesson based on the teacher’s plans. This may explain that the brightness levels differed between the different instructional activities due to varying ambient lightning conditions from varying cloud cover across the lesson, rather than from the instructional activities themselves. To investigate this, a windowless classroom study would need to be performed.

One technical limitation of our study is that the AI estimated movement and the normalised brightness change variables correlated significantly (See Extended Data Table 1) which could have affected the results of the multiple regression modelling due to multicollinearity. In this model movement was not identified as a significant predictor of the composite phasic sympathetic activation. Although a VIF below 1.3 suggests no multicollinearity (Lavery et al., 2019), the associated correlation might still lead to a Type II error, whereby an existing effect of movement on sympathetic activation is missed. Furthermore, the OpenPose AI detection cannot identify body parts behind desks and may not have caught all body movement. As such, more research into the effect of movement on tonic and phasic sympathetic activation in learning situations should be conducted.

Another limitation of the current study is that even though the results regarding the connection between phasic sympathetic activation and learning was replicated across two contexts, the range of the linear relationship remains unclear. Further studies should investigate these limitations.

Despite these limitations, our study contributes to the field by introducing a candidate for a physiological indicator for learning which is affected by sensory input and instructional activities and operates on a timescale appropriate for use in learning technology development and learning intervention assessment.

## Methods

The study was approved by the south-east Regional Ethics Committee (ref. 640037) and the Norwegian Centre for Research Data (ref. 489494) and students (and their guardians if they were below 16 years) provided informed consent for participation. We gave each participant a unique and anonymous ID. If not all students had agreed to the video data collection, the video camera would have been angled in such a way that the students not agreeing to the video data collection would be outside the angle of recording.

### 1. Participants and context

#### 1.1 Study one

The data was collected in a physics classroom in vocational education with 12 students (17-18 years, all male) and one teacher. SC was collected for all students, as was the video data. Video data was collected from one camera situated at the front of the classroom facing the students. The teacher was included in the video data when not standing in the front of the classroom but moving among the students. The data was collected during one 135-minute lesson, excluding two 10-minute breaks and the fastening of the SC sensors on the students, which resulted in 75 minutes of synchronised video and SC data. The teacher went through solutions to physics tasks whilst interacting with the students, which the students had worked on previously. Afterwards, the students worked further on the same or similar physics tasks while the teacher helped the students.

The study included a post-test design, where students were tested in physics according to the guidelines of the European Union Aviation Safety Agency (EASA). This knowledge test consisted of multiple-choice questions with three alternatives, with only one correct alternative. The test was given on paper and consisted of 40 questions. Example questions can be found in Figure 1, whilst the complete test is confidential due to EASA regulations (EASA, n.d.).

**Fig. 1.**
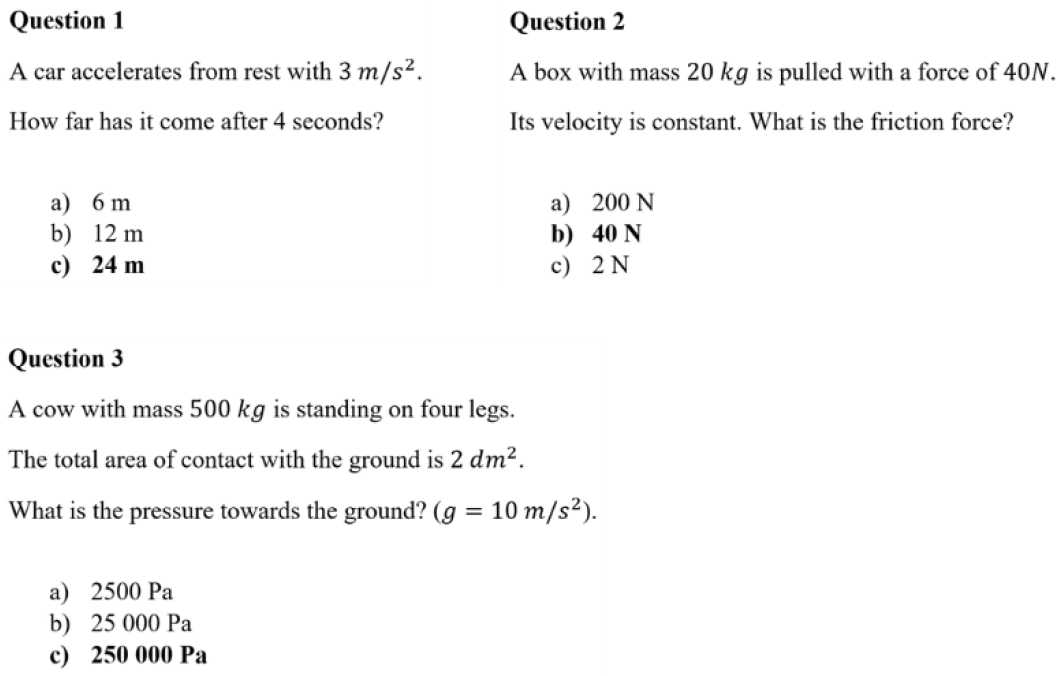
Examples of test questions in the classroom study. The correct answers are in bold.

#### 1.2 Study two

The data was collected at a cognitive lab at a Norwegian university with 100 participants, where 59 were self-reported female, 39 were self-reported male, and two preferred not to disclose. Participants were aged 12 – 21 years (M = 16.52, SD = 1.73) and were recruited through their schools sending out information about the project. Exclusion criteria were a diagnose of ME/CFS, ADHD, autism specter disorders, dyslexia, any learning disabilities, depression, anxiety or similar diagnoses or disabilities.

Before the main data collection, participants answered a digital survey of their spatial abilities as spatial intelligence are considered central for learning in STEM (science, technology, engineering, and mathematics) (Wai et al., 2009).

The Spatial Reasoning Instrument (SRI) is designed to assess spatial reasoning abilities in children and youth by evaluating four distinct aspects of spatial ability: mental rotation (the ability to mentally rotate 2D and 3D objects), spatial visualization (the ability to mentally visualize 3D objects), spatial orientation (the mental capability to orient oneself in space), and spatial relations (which encompass the relationships among the previous three aspects) (Ramful et al., 2017). This instrument demonstrates strong psychometric properties regarding validity and reliability (Ramful et al., 2017). A Norwegian translation of the SRI has been validated with a sample of 535 Norwegian students aged 9 to 16 (Flø & Smedsrud, 2025).

The lab data collection lasted about 50 minutes and was run at the university lab with one participant and one data collection supervisor per session. During the data collection, the participant was seated in a comfortable chair with an approximate distance of 60 cm between the computer monitor and their eyes. This data collection consisted of one pretest at the lab when one of the data collection supervisors set up the equipment to record skin conductance.

When the SC electrodes were in place, the participants performed two relaxation tasks (i.e., watching a one-minute relaxing nature video, and 14 minutes of hearing clicks with eyes closed in a sensory gating paradigm), and two cognitively demanding tasks (i.e., watching a six-minute instructional video, and a 2-back test). Only SC data from watching the instructional video will be reported on here. The instructional video was randomly shown in one of two versions, where colour saturation and viewing volume had been altered between them, to provide different amounts of sensory stimuli. After performing the tasks, the participants answered the post-test and were paid 500 NOK (∼ 45 €).

The study included a pre-/post-test design for the biology knowledge test. The knowledge test consisted of 10 multiple-choice questions with four alternatives, with only one correct alternative. The tests were given digitally, and to measure potential learning, the same tests were given before and after the series of workshops. The complete test can be found in Figure 2.

**Fig. 2.**
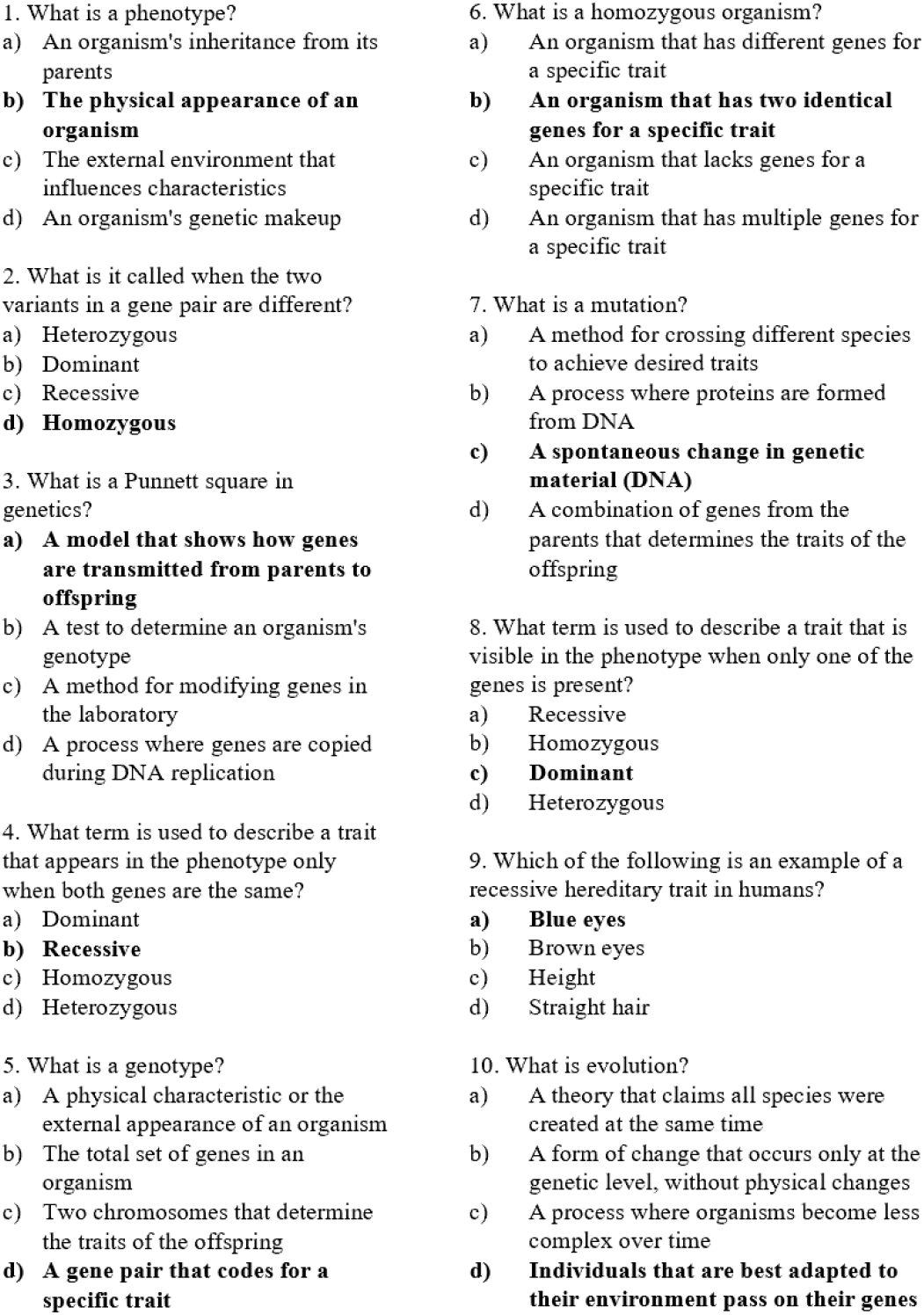
The learning test used in the lab study. The correct answers are in bold.

### 2. SC measurement and analysis

#### 2.1 Measurement

Because there are wearable instrument validity issues for SC measurement (Hu et al., 2024; Ronca et al., 2023), we have chosen to use the BITalino SC sensor in study one, which has been sufficiently validated against laboratory measures (Batista et al., 2019). SC was measured at 10 Hz with two solid-gel self-adhesive disposable Ag/AgCl electrodes fastened to the students second and third proximal phalanges of the non-dominant hand, exceeding the suggested 5 Hz minimum sampling rate for spontaneous SC responses (Tronstad et al., 2022). The classroom temperature was maintained at 22 °C to prevent any thermoregulatory interference with the SC measurement (Boucsein, 2012) and was the standard setting for the thermostat in the school.

In the cognitive lab in study two, we collected SC data with a Brainproducts ActiChamp Plus system with a BP-MB-30 GSR module incl. electrodes and gel at 5000 Hz with the same electrode placement and temperature setting as in study one. The electrodes were carefully taped to the skin with medical tape.

#### 2.2 Analysis

We used the MATLAB based software Ledalab (version 3.4.9; http://www.ledalab.de/) for SC signal processing. Prior to analysis, we performed a visual inspection for identification of motion artefacts according to decision rules from previous work (Kleckner et al., 2017). Data from one person in the lab study was excluded on this basis, and no data from the classroom study. There was no SC data for another person in the lab study due to technical issues, bringing the total number of participants to 98.

The raw signal serves as the default input for Ledalab software in conducting the decomposition analysis of the SC signal (Benedek & Kaernbach, 2010), i.e., no preprocessing of the SC signal was performed. As the recommended method for decomposition of the SC signal into tonic (slow-varying) and phasic (fast-varying) components is the continuous decomposition analysis (Benedek & Kaernbach, 2010), this algorithm was used in both study one and study two. In study two, SC data was down-sampled to 10 Hz prior to analysis as sampling rates of 5 Hz will preserve the relevant signal information, especially for long-term recordings of spontaneous SC (Tronstad et al., 2022), whilst in study one the SC data was collected at 10 Hz, leaving no need for down-sampling.

The tonic component is often considered a baseline signal whereas the phasic component is considered a more immediate and reactive response, often used to assess immediate responses in arousal (Villanueva et al., 2018). Because baselines are important to correct physiological data which can vary according to e.g., person or time of day (Cowley et al., 2014), we chose to mainly focus on the phasic component of the SC signal. Moreover, because there is a response latency of ∼1s (Sjouwerman & Lonsdorf, 2019; Society for Psychophysiological Research Ad Hoc Committee on Electrodermal Measures, 2012), we used one-minute segments for the analyses.

### 3. Video analysis

The video analysis was performed similarly for both studies with the exception of excluding the AI estimated movement for study two as the participants watched the video, and did not participate in it, so that the AI movement would reflect sensory input as already measured by the normalised brightness change variable. Additionally, instructional methods were not assessed for study two because the whole instructional method was defined as a “learning film”.

#### 3.1 Instructional methods (qualitative)

The two authors independently identified instances of the five different instructional methods in the 75-minute video material. We categorised one-minute segments according to the instructional design that was dominant, i.e., present for more than half of the minute in question. Five minutes were omitted as they were mainly practical messages to the students. Interrater reliability for the five categories was 0.92 (Cohen’s kappa). Any preliminary disagreements were discussed and resolved to complete agreement.

#### 3.2 Brightness

Brightness is a perceptual attribute of the human visual system and must be calculated, not physically measured (Judd, 1952). Video and browsers usually use the sRGB colour space to represent the brightness values of each pixel which needs to be transformed to the RGB colour space by an inverse gamma function to linearly add the values for each colour. This grey-scale conversion was done by processing each frame by converting the RGB values of pixels to a grey-scale brightness value using predefined luminance coefficients (R = 0.212655, G = 0.715158, B = 0.072187) (International Telecommunication Union, 1990) and an inverse gamma correction function, approximated by exponentiation by 1/(2.2) (to avoid numerical issues close to zero). The following intensity computation summarises the grey-scale brightness for each frame and this value is again summarised for each one-minute segment of video.

#### 3.3 Normalised brightness change

As mammals have velocity sensitive visual neurons (Orban et al., 1981; Orban et al., 1986), we included change in brightness as a sensory input variable. This may reflect movement from the participants as well as their sensory input in the classroom study (study one) but does not include movement in the lab study (study two) as the video analysis was performed on the learning video the participants watched. To investigate a potential effect of student movement more directly in study one where the students were part of the analysed video, an AI movement variable was included (see the 3.5 AI movement section). Moreover, to enable comparison across different brightness conditions, the variable was normalised by dividing the change in brightness by the original brightness level. The resulting definition for normalised brightness change is the difference in brightness between two consecutive frames of video relative to the brightness of the first frame, summarised for each one-minute section of video.

#### 3.4 Loudness

To account for the way human hearing works, we used the loudness method in the pyloudnorm python library which calculates the loudness of an audio signal based on the EBU R128 loudness standard (International Electrotechnical Commission, 2003). The process involves filtering (with a K-weighting filter), calculation of the absolute squared value of the filtered signal, time-averaging using gating to find momentary loudness, and computing an overall loudness measurement over the entire audio signal expressed in Loudness Units relative to Full Scale (LUFS).

The gating technique implemented in pyloudnorm, is designed to ignore very quiet parts of the audio that wouldn’t significantly contribute to the perceived loudness, as human perception of loudness is more sensitive to louder sounds (Uppenkamp & Röhl, 2014). A graphical overview of the steps of the loudness calculation can be found in Figure 3.

**Fig. 3.**
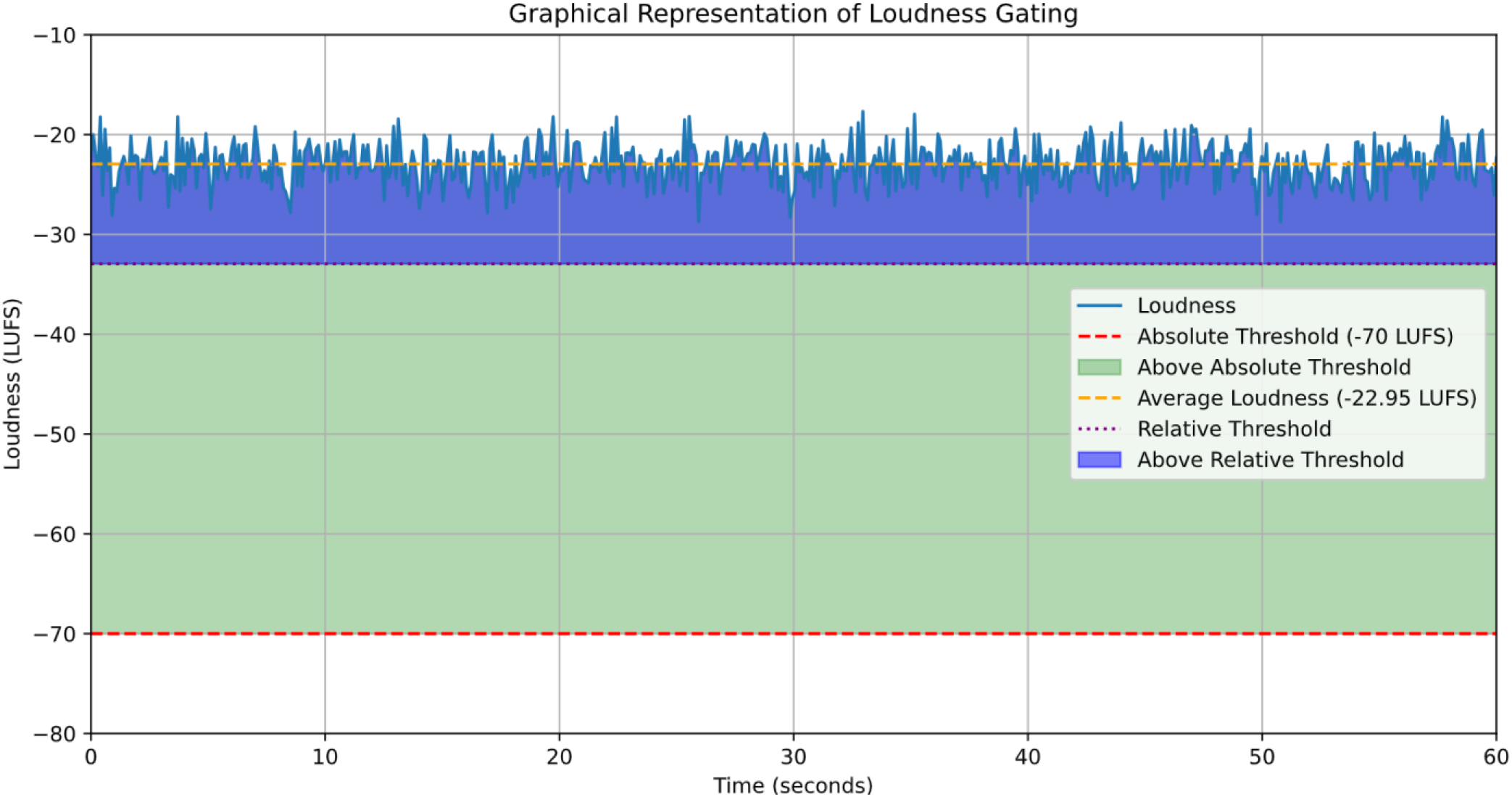
Graphical representation of loudness gating. Loudness measured by LUFS is the part of the sound signal above an absolute and relative threshold and was calculated using the loudness method in the pyloudnorm python library which is in accordance with EBU R128 loudness standard.

### 3.5 AI estimated movement

An additional analysis of movement using the OpenPose AI model (https://github.com/CMU-Perceptual-Computing-Lab/openpose) for pose estimation was performed. We used the same 1 Hz compressed one-minute segments as in the brightness and loudness analysis, but all pixels were kept for each frame. The OpenPose AI model returns one JSON-file per frame (i.e., 60 per minute in this study) containing the coordinates of 25 body parts for each identified person in the frame. To translate these coordinates to movement, the individual persons need to be traced across frames, which was done by using software connecting the person closest to where the person was in one frame to the next frame (Hur & Bosch, 2022). This tracing was done within each minute segment. After this step, self-developed software (GitHub-link removed for anonymised review) was used to calculate the distance between each of the 25 body parts belonging to one person across the 60 frames in a minute and doing so for all persons in the one-minute segment. There was no need to normalise across student number, as all students were present in each frame. Thus, the AI-based analysis estimate movement only based on people, not other objects which allow object movement to be controlled for.

### 4. Statistical analyses

#### 4.1 Correlation and multiple regressions

We used R version 4.3.1 (R Core Team, 2023) with R studio version 2023.12.1 for the statistical analysis and estimated the IRT models with the R package mirt (Chalmers, 2012).

Prior to the multiple regressions, we checked the pairwise correlation between the variables to identify potential multicollinearity issues. Potential issues with high correlations are the reduced estimates of the correlation coefficients and inflation of type 2 error (Lavery et al., 2019). Suggested remedies have been excluding variables or combining them (Franke, 2010). Because the two phasic SC variables demonstrated a significant correlation of r = 0.92, we created a new phasic SC variable to preserve the relationships between both of the original phasic SC variables and the sensory input and AI movement variables, resulting in the new composite phasic variable, SC_Phasic_M_/SC_Phasic_SD_.

Then, data from study two was analyses through a multiple regression procedure to investigate the relationship between learning outcome (i.e., the difference between post-test and pretest scores) and the SC_Phasic_M_/SC_Phasic_SD_ variable, controlling for gender and spatial ability. All variables were standardised to enable comparison between effect sizes. Afterwards, to assess whether sensory input or movement could predict the phasic SC variable another multiple regression was performed with all variables standardised.

Both multiple regression models were checked for the normality of residuals by visual inspection of their Q-Q-Plots and the Shapiro-Wilk test which indicated normality. The Variance Inflation Factor (VIF) for each predictor in both models were all below 2, dispelling multicollinearity concerns.

#### 4.2 Item response theory (IRT)

The spatial ability scores were estimated using a validated IRT based model from a data set of 535 children representative of the general population (Flø & Smedsrud, 2025) with standardised scores. It is an underlying assumption of IRT modelling that the estimated trait is normally distributed (Lord, 1986), which the population sample used for model development satisfied (Flø & Smedsrud, 2025). This model is based on logarithmic models for the probability of each of the test items being correct based on the student’s ability level (Chalmers, 2012). The estimated reliability of the EAP (expected a posteriori) ability estimates was calculated (Bock & Mislevy, 1982), resulting in an acceptable value of 0.83 (Maydeu-Olivares, 2013), the IRT reliability estimate corresponding to the classical Cronbach’s alpha.

#### 4.3 Group differences

To assess group differences, the data were first checked for normality using the Shapiro-Wilk test. As no variables were normally distributed, the parametric Mann-Whitney U test for unpaired data was used to check for significant group differences with the corresponding effect size measure, namely the rank biserial correlation, r (Tomczak & Tomczak, 2014).

## Data availability

The video dataset generated and analysed during the current study are not publicly available due to ethical restrictions and participant confidentiality agreements, whilst the rest of the data are available from the corresponding author upon reasonable request.

## Code availability

All code used for data analysis is available at GitHub - gmfloe/Video-analysis.

