## Extended data for "Sympathetic activation of sensory input and learning"

Table 1. Correlations for all sensory input, movement, and SC variables per minute for the classroom setting in study one.

| | *Brightness* | *Normalised brightness change* | *Loudness* | *AI-movement* | $SC_{tonic\_M}$ | $SC_{tonic\_SD}$ | $SC_{phasic\_M}$ | $SC_{phasic\_SD}$ |
|---|---|---|---|---|---|---|---|---|
| Normalised brightness change | -0.06 | | | | | | | |
| Loudness | 0.008 | **-0.33**** | | | | | | |
| AI-movement | **-0.24*** | **0.32**** | -0.21 | | | | | |
| $SC_{tonic\_M}$ | 0.10 | 0.12 | **0.38***** | 0.10 | | | | |
| $SC_{tonic\_SD}$ | **0.29*** | **0.28*** | 0.08 | 0.15 | **0.44***** | | | |
| $SC_{phasic\_M}$ | 0.20 | 0.18 | **0.30**** | 0.05 | **0.54***** | **0.76***** | | |
| $SC_{phasic\_SD}$ | **0.34**** | 0.12 | 0.19 | 0.05 | **0.46***** | **0.80***** | **0.92***** | |
| $SC_{phasic\_M}$/ $SC_{phasic\_SD}$ | **-0.34**** | 0.10 | **0.30**** | -0.05 | 0.08 | -0.19 | 0.07 | **-0.30**** |

* denotes p-values < 0.05, ** denotes p-values < 0.01, and *** denotes p-values <0 .001

Tonic and phasic SC

Table 2. Standardised regression coefficients for the multiple regression for tonic SC as outcome variable.

| | *Estimate, β* | *SE* | *t-value* | *p-value* |
|---|---|---|---|---|
| Brightness | 0.15 | 0.11 | 1.43 | 0.16 |
| Normalised brightness change | 0.24 | 0.11 | 2.05 | **0.044*** |
| Loudness | 0.49 | 0.11 | 4.39 | **4.0e-5***** |
| AI-movement | 0.16 | 0.11 | 1.44 | 0.15 |

* denotes p-values < .05, ** denotes p-values < .01, and *** denotes p-values < .001

Adjusted R-squared = 0.19 (Wherry1), F(4,70) = 5.62 with corresponding **p-value = 0.0005**.

Table 3. Standardised regression coefficients for the multiple regression for phasic SC as outcome variable.

| | *Estimate, β* | *SE* | *t-value* | *p-value* |
|---|---|---|---|---|
| Brightness | 0.24 | 0.11 | 2.20 | **0.031*** |
| Normalised brightness change | 0.30 | 0.12 | 2.64 | **0.010*** |
| Loudness | 0.42 | 0.11 | 3.76 | **0.00034***** |
| AI-movement | 0.092 | 0.11 | 0.81 | 0.42 |

* denotes p-values < .05, ** denotes p-values < .01, and *** denotes p-values < .001

Adjusted R-squared = 0.18 (Wherry1), F(4,70) = 5.33 with corresponding **p-value = 0.0008**.

### Brightness

Table 4. Descriptive statistics for the brightness levels for the different instructional activities.

| *Instructional activity* | *Brightness M* | *Brightness SD* |
|---|---|---|
| Blackboard w/questions | 644.57 | 25.87 |
| Teacher-led discussion | 640.63 | 17.43 |
| Tasks with help | 644.76 | 16.36 |
| Tasks without help | 639.03 | 23.97 |
| Blackboard wo/questions | 642.88 | 37.53 |

Table 5. The group comparisons for the brightness levels for the different instructional activities in the classroom study.

| *Instructional activity* | *Instructional activity* | *Test statistic, W* | *Effect size* | *Conf. low* | *Conf., high* | *p-value* |
|---|---|---|---|---|---|---|
| Blackboard w/questions | Blackboard wo/questions | 193451 | 0.06 | 0.02 | 0.10 | **0.006**** |
| Blackboard w/questions | Teacher-led discussion | 672512 | 0.05 | 0.01 | 0.09 | **0.009**** |
| Blackboard w/questions | Working tasks with help | 313929 | 0.05 | 0.02 | 0.09 | **0.01*** |
| Blackboard w/questions | Working tasks without help | 1087980 | 0.05 | 0.01 | 0.08 | **0.008**** |
| Blackboard wo/questions | Teacher-led discussion | 54666 | 0.06 | 0.006 | 0.12 | 0.11 |
| Blackboard wo/questions | Working tasks with help | 23948 | 0.21 | 0.13 | 0.29 | **9.41e-07***** |
| Blackboard wo/questions | Working tasks without help | 88403 | 0.05 | 0.008 | 0.1 | 0.06 |
| Teacher-led discussion | Working tasks with help | 137529 | 0.14 | 0.08 | 0.19 | **1.8e-05***** |
| Teacher-led discussion | Working tasks without help | 356355 | 0.004 | 0.0006 | 0.05 | 0.86 |
| Working tasks with help | Working tasks without help | 215394 | 0.09 | 0.04 | 0.13 | **0.001**** |

* denotes p-values < 0.05, ** denotes p-values < 0.01, and *** denotes p-values < 0.001

### Normalised brightness change

Table 6. Descriptive statistics for the normalised brightness change levels for the different instructional activities.

| *Instructional activity* | *Normalised brightness change M* | *Normalised brightness change SD* |
|---|---|---|
| Blackboard w/questions | 28.07 | 14.41 |
| Teacher-led discussion | 30.09 | 10.76 |
| Tasks with help | 43.94 | 16.50 |
| Tasks without help | 44.67 | 16.13 |
| Blackboard wo/questions | 25.16 | 12.38 |

Table 7. The group comparisons for the normalised brightness change levels for the different instructional activities in the classroom study.

| *Instructional activity* | *Instructional activity* | *Test statistic, W* | *Effect size* | *Conf. low* | *Conf., high* | *p-value* |
|---|---|---|---|---|---|---|
| Blackboard w/questions | Blackboard wo/questions | 238164 | 0.19 | 0.15 | 0.23 | **2.2e-16*** ** |
| Blackboard w/questions | Teacher-led discussion | 569756 | 0.07 | 0.03 | 0.11 | **0.0003*** ** |
| Blackboard w/questions | Working tasks with help | 216576 | 0.23 | 0.19 | 0.27 | **2.2e-16*** ** |
| Blackboard w/questions | Working tasks without help | 680532 | 0.28 | 0.25 | 0.31 | **2.2e-16*** ** |
| Blackboard wo/questions | Teacher-led discussion | 31946 | 0.33 | 0.26 | 0.39 | **2.2e-16*** ** |
| Blackboard wo/questions | Working tasks with help | 10762 | 0.54 | 0.49 | 0.6 | **2.2e-16*** ** |
| Blackboard wo/questions | Working tasks without help | 31698 | 0.41 | 0.36 | 0.45 | **2.2e-16*** ** |
| Teacher-led discussion | Working tasks with help | 151278 | 0.23 | 0.17 | 0.29 | **1.8e-13*** ** |
| Teacher-led discussion | Working tasks without help | 438504 | 0.20 | 0.15 | 0.25 | **2.2e-16*** ** |
| Working tasks with help | Working tasks without help | 206402 | 0.05 | 0.004 | 0.11 | 0.05 |

* denotes p-values < 0.05, ** denotes p-values < 0.01, and *** denotes p-values < 0.001

### Loudness

Table 8. Descriptive statistics for the loudness levels measured in LUFS for the different instructional activities.

| *Instructional activity* | *Loudness M* | *Loudness SD* |
|---|---|---|
| Blackboard w/questions | 509.45 | 1200.94 |
| Teacher-led discussion | 542.61 | 623.32 |
| Tasks with help | 820.45 | 800.59 |
| Tasks without help | 653.10 | 569.69 |
| Blackboard wo/questions | 307.13 | 262.88 |

Table 9. The group comparisons for the loudness levels for the different instructional activities in the classroom study.

| *Instructional activity* | *Instructional activity* | *Test statistic, W* | *Effect size* | *Conf. low* | *Conf., high* | *p-value* |
|---|---|---|---|---|---|---|
| Blackboard w/questions | Blackboard wo/questions | 179727 | 0.02 | 0.0009 | 0.06 | 0.32 |
| Blackboard w/questions | Teacher-led discussion | 701443 | 0.09 | 0.05 | 0.12 | **1.2e-05*** ** |
| Blackboard w/questions | Working tasks with help | 545693 | 0.37 | 0.34 | 0.41 | **2.2e-16*** ** |
| Blackboard w/questions | Working tasks without help | 1400550 | 0.30 | 0.27 | 0.33 | **2.2e-16*** ** |
| Blackboard wo/questions | Teacher-led discussion | 64469 | 0.06 | 0.006 | 0.12 | 0.07 |
| Blackboard wo/questions | Working tasks with help | 51589 | 0.49 | 0.42 | 0.55 | **2.2e-16*** ** |
| Blackboard wo/questions | Working tasks without help | 132741 | 0.23 | 0.17 | 0.28 | **1.6e-15*** ** |
| Teacher-led discussion | Working tasks with help | 57362 | 0.43 | 0.38 | 0.48 | **2.2e-16*** ** |
| Teacher-led discussion | Working tasks without help | 270118 | 0.20 | 0.15 | 0.25 | **2.2e-16*** ** |
| Working tasks with help | Working tasks without help | 120776 | 0.28 | 0.23 | 0.34 | **2.2e-16*** ** |

* denotes p-values < 0.05, ** denotes p-values < 0.01, and *** denotes p-values < 0.001

### AI estimated movement

Table 10. Descriptive statistics for the AI estimated movement levels for the different instructional activities.

| *Instructional activity* | *Movement M* | *Movement SD* |
|---|---|---|
| Blackboard w/questions | 571.71 | 743.83 |
| Teacher-led discussion | 560.99 | 717.94 |
| Tasks with help | 647.44 | 747.25 |
| Tasks without help | 656.89 | 863.87 |
| Blackboard wo/questions | 394.37 | 287.61 |

Table 11. The group comparisons for the AI estimated movement levels for the different instructional activities in the classroom study.

| *Instructional activity* | *Instructional activity* | *Test statistic, W* | *Effect size* | *Conf. low* | *Conf., high* | *p-value* |
|---|---|---|---|---|---|---|
| Blackboard w/questions | Blackboard wo/questions | 185328 | 0.04 | 0.008 | 0.08 | 0.052 |
| Blackboard w/questions | Teacher-led discussion | 617824 | 0.02 | 0.0009 | 0.05 | 0.44 |
| Blackboard w/questions | Working tasks with help | 309150 | 0.06 | 0.02 | 0.11 | **0.003**** |
| Blackboard w/questions | Working tasks without help | 832243 | 0.16 | 0.12 | 0.20 | **2.2e-16***** |
| Blackboard wo/questions | Teacher-led discussion | 52072 | 0.08 | 0.01 | 0.14 | 0.02 |
| Blackboard wo/questions | Working tasks with help | 25631 | 0.16 | 0.08 | 0.24 | **0.0002***** |
| Blackboard wo/questions | Working tasks without help | 66819 | 0.18 | 0.13 | 0.23 | **1.0e-10***** |
| Teacher-led discussion | Working tasks with help | 128135 | 0.07 | 0.01 | 0.13 | **0.03*** |
| Teacher-led discussion | Working tasks without help | 416344 | 0.15 | 0.10 | 0.19 | **1.0e-09***** |
| Working tasks with help | Working tasks without help | 177768 | 0.06 | 0.01 | 0.11 | **0.02*** |

* denotes p-values < 0.05, ** denotes p-values < 0.01, and *** denotes p-values < 0.001
